# Enhancements in Squid Retinal Responses to Change of Polarizations in a Caustic Shallow Water

**DOI:** 10.1101/2024.04.17.589948

**Authors:** Jing Cai, Sergei Nikonov, Alison M. Sweeney

## Abstract

Marine animals with polarization vision are able to effectively detect moving objects in shallow waters, which are illuminated by dynamic fluctuations of downwelling light known as caustics. While behavioral studies across different animal species have demonstrated the support of polarization vision in moving object detection within this noisy environment, little is known about how their retinal photoreceptors, absorbing polarized photons, respond to moving objects, or how each photoreceptor contributes to the collective retinal reaction to changes in polarization. In this study, we employed multi-electrode array recordings to examine the retinal neural response of squid to polarized light stimuli that were designed to simulate caustics environment. Extracellular retinal recordings not only exhibit neural activities selective to the direction of polarization but also demonstrate a significant enhancement in response to stimuli with changing polarization compared to constant polarization. This enhancement is robust in almost all recording channels, but absent in a random permutation of the recordings from different trial types. These results suggest that the retinal photoreceptors directly encode the change of polarization stimuli, thereby contributing to signal detections with polarization vision. Together, our research represents a novel neural exploration of cephalopod polarization vision in a caustic environment, and advances our understanding of how nature parses scenes with salient, dynamic polarization in animal vision.

## Introduction

In shallow water environments, downwelling light can refract from surface ripples, creating dynamic bands of intense illumination known as caustics (*1-3*). This visual noise may help conceal the movement of animals (*4*), making it challenging to detect visual cues from predators and prey (*5*). Concurrently, the background downwelling light is partially linearly polarized (*6*), thereby generating polarization contrasts when reflecting from the surfaces of marine animals (*7, 8*). The change of polarization contrasts from a moving object in a noisy environment may provide a perceptual opportunity for object detection (*9, 10*). Indeed, behavioral experiments on animals with polarization vision demonstrate that dynamic caustics do hinder the detection of unpolarized stimuli, but not the detection of polarized stimuli (*11, 12*). Therefore, we hypothesize that polarization vision assists in movement detection by responding to changes of polarizations under the caustic environment.

Behavioral experiments demonstrate that cephalopods perceive polarization (*9, 13, 14*). The ultrastructure of the retina suggests a mechanism for differential absorption of different orientations of linear polarization (*15-18*). Each retinal cell has highly oriented, parallel arrays of microvilli, and the retina is everted, with the photoreceptor cell layer close to the lens in the camera-type eye (*19-21*). This anatomy allows for the precise alignment of rhodopsin molecules along well-defined axes of microvilli, in turn ensuring each photoreceptor interacts solely with specific polarizations (*22, 23*). The size of these retinal photoreceptors typically ranges from 2 to 5 microns in cross-section, with the estimated angular acceptance of ∼ 10^−4^ rad. On the proximal side of the photoreceptors, action potentials are output from an axon that projects directly to the optic lobe (*24, 25*). Across the retina, photoreceptors are organized so that adjacent cells absorb perpendicular polarizations (*23, 24*). Decades ago, studies of single-neuronal recordings showed tuning curves of action potentials with rotating polarization orientation, confirming the ability of individual retinal cells to respond to light of a specific polarization orientation (*26*). However, little is known about how populations of photoreceptors collectively respond to polarization in a scene, or how the organization of the retina allows for perception of moving objects under caustic illumination conditions.

In this study, we aim to gain insight into how the squid retina responds to scenes with moving reflective objects in dynamic, caustic illumination using a simplified experimental paradigm. We employed multi-electrode arrays to simultaneously record neural activity from dozens of channels across the area of a single squid retina. For stimuli, we used full-field flashes with randomized durations and intervals to simulate caustic flickers, using a linearly polarized light source to mimic reflection from a target object. In some trials, the orientation of the polarized stimulus was rotated between flashes to mimic the movement of a polarized target object. This set of stimuli provided a simplified scenario for probing the detection of moving objects amidst the background light fluctuations caused by caustics. We first examined the extra-cellular neuronal responses to the onset of illuminations. We then performed statistical analysis to study the retinal responses under the condition representing the movement of a target as compared to a linearly polarized background illumination. Our results show a significant enhancement in the response of retinal cells to changes in polarization, and this effect was observed consistently across individual channels. This finding suggests that the squid retina possesses a heightened sensitivity to changes of polarized scenes, indicating their integral role in facilitating signal detection in polarization perception.

## Material and Methods

### Animal and preparation

Specimens of the squid *Doryteuthis pealeii* were obtained freshly decapitated from the Marine Biological Laboratory at Woods Hole, MA. A fresh eye was dissected in a petri-dish containing micron-filtered seawater. Optic fibers on the posterior retina were carefully removed, and the retina was carefully dissected from remaining eye tissues. A 5-mm diameter biopsy punch was then used to excise a central retinal sample. All the preparations were performed on ice under room light.

### Experimental paradigm

To test how squid photoreceptors respond to moving stimuli of approaching predators or prey, or signals from conspecifics in shallow waters with dynamic caustics, we designed experimental trials with illuminations of changing polarization orientation. Specifically, we used full-field illumination events with randomized durations and intervals to represent the flickers from caustics. During the interval, we either rotated the polarization by 90 degrees in the same plane as the retina to represent a movement of the object, or kept the polarization orientation fixed. Although this paradigm does not exactly replicate the natural conditions we aim to examine, it provides a simplified representation of such conditions over ecologically relevant timescales.

Simply put, our experiments included two sets of trials. One set tested responses to calibrated flashes of light with the direction of polarization changed between flashes, and the other set tested responses to light of fixed polarization. The alternation trials involved a switch between the two polarizations during every dark interval, whereas the polarizer was fixed in one orientation for constant polarization trials. The orientation of the polarizer was chosen arbitrarily with respect to the orientation of the retina, and in the trials in which polarization changed, it was rotated 90º from this initial orientation. The duration of each trial was 150 seconds, with the flash timing manually controlled.

### Multi-electrode array (MEA) recording

To investigate the response of retinal activities to different polarization states, we employed a multi-electrode array (MEA) for simultaneous recording at multiple locations on the retina. The isolated retina was carefully positioned in the recording chamber with the photoreceptor-side of the retina in contact with the array. This configuration mimicked the natural light path in an intact eye, ensuring that light stimuli first reached the photoreceptors (**Figure 1B**). To improve retinal contact with the electrodes, an anchor (0.8 g, HSG-MEA-5BD, ALA Scientific, Farmingdale, NY, USA) was placed on top of the retinal tissue (**Figure 1A, B**). The recording chamber was perfused with filtered seawater with the temperature maintained at 10 °C during recording.

**Figure 1.**
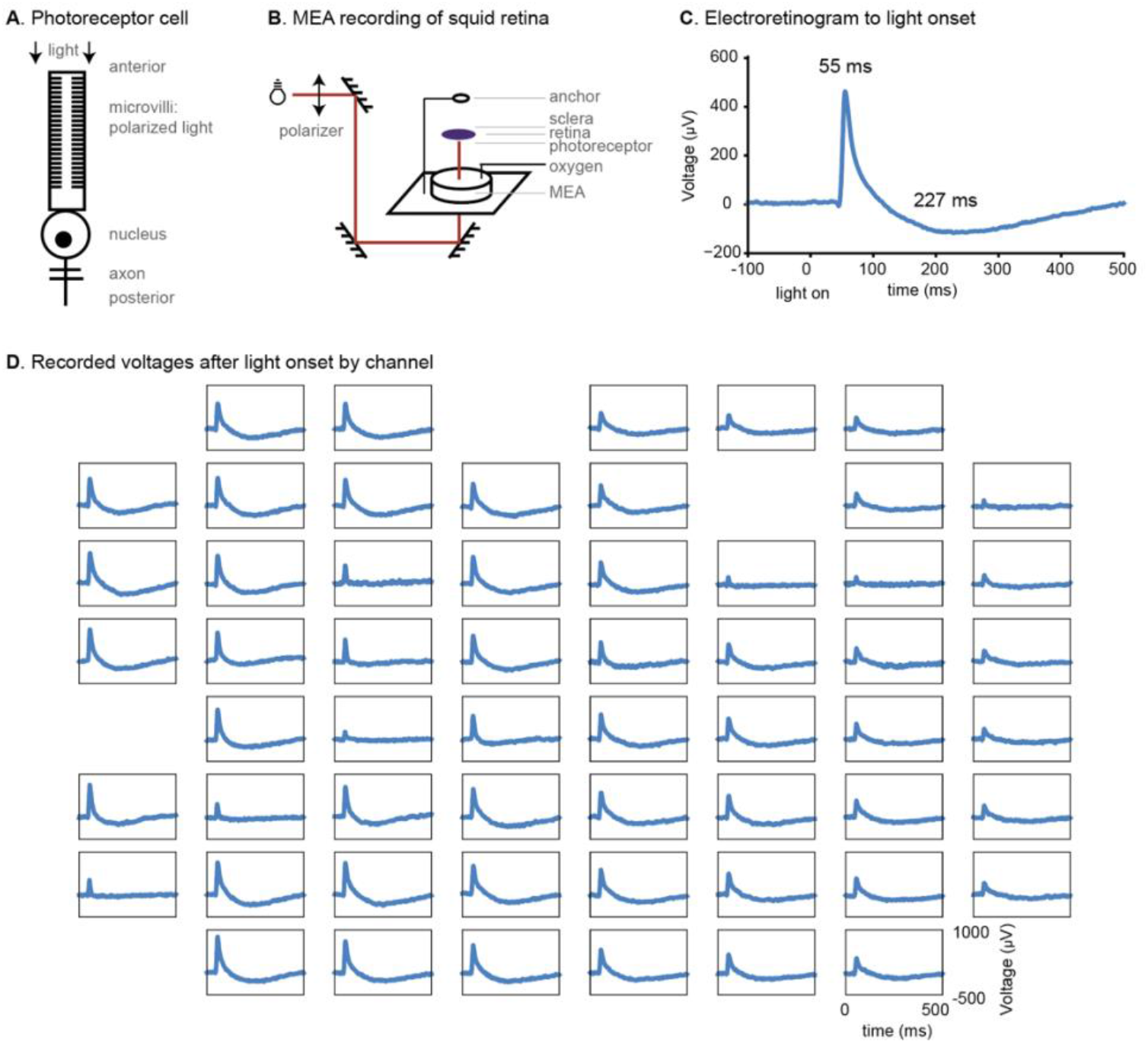
Multi-electrode array recordings on squid retina. **A**. Illustration of the structure of a cephalopod retinal cell. **B**. Experimental setup for multi-electrode array (MEA) recordings on squid retinal tissue. Polarized incident light (*dark red*) was reflected to the photoreceptor side of the retinal tissue with their extra-cellular voltages recorded by MEA. **C**. On average, retinal cells responded to the onset of illumination with an initial maximum at 55 milliseconds, and a refractory minimum at 227 milliseconds. **D**. Retinal response to onset of illumination across channels that exhibited significant signal-to-noise ratio.

Extra-cellular voltages of the retina were measured by 60 electrodes in the standard arrays (60MEA200/30IR–ITO–G). Each electrode is 30 μm in diameter with a 200 μm inter-electrode distance (Multichannel Systems, Reutlingen, Germany). Therefore, given the characteristic size of a squid photoreceptor cell, a single channel potentially recorded voltages from a small cluster of neurons, while different channels were likely to record distinct clusters of neurons. The raw voltages were recorded by an MEA amplifier (MEA1060-Inv, Multichannel Systems, Reutlingen, Germany) and then digitized at 10 KHz by custom-made acquisition software in LabView (National Instruments, Austin, TX, USA).

Meanwhile, the light stimuli were recorded on a separate analog channel simultaneously with the neural recordings. Next, all recorded data were combined to a single dataset in MATLAB (MathWorks, Natick, MA, USA).

The light stimuli were generated with a full-field 455 nm light source with intensity on the order of 10^2^ W m^-2^ (455 nm 75 lm Diamond Dragon series LED, Osram Opto Semiconductors/Digi-Key, Thief River Falls, MN). The light source was driven by Clampex software and digitizer (Digidata 1440, Molecular Devices, San Jose, CA, USA). Calibrated full-field light passed through a linear polarizing filter and was delivered by a 4x microscope objective lens onto the retina.

### Retinal responses to constant polarization and alternation of polarization

We investigated how photoreceptors recorded by different channels responded to these stimuli. To start, we selected channels that showed significant responses to the onset of illumination. We averaged these cell activities across all trials and events. We then compared the maximum of voltages within the first 300 milliseconds after light onset to the voltages during the interval of 100 to 500 milliseconds before light onset. We defined as responsive any channel in which the maximum voltage after light-on was six standard deviations greater than the voltage before light-on. These responsive channels were selected for further analysis.

Next, we examined whether any of the responsive channels showed a different response to one polarization orientation relative to the perpendicular counterpart. Because the absolute voltages of neural responses depended on the duration of the dark interval before a flash, a direct comparison between the two conditions may result in false positives due to the variance of the dark duration. Therefore, we binned the neural responses based on the duration of the dark interval for each channel, and calculated the voltage difference between the two polarization conditions within each bin. After this process, we performed a T-test to examine whether the differences of polarization were significantly different from zero. Channels with resulting p-values less than 0.05 were identified as polarization-selective channels.

The latency of the extrema were not dependent on the durations of dark interval. Therefore, we simply combined all observations for each polarization without categorizing by interval durations, and used a two-sided T-test to determine whether a channel showed significant differences between the two constant polarizations. After these procedures, we performed a permutation test to evaluate whether the fraction of channels showing a significant difference in polarization response exceeded what could be attributed to chance. To estimate the fraction of responses that could be due to random fluctuations, we combined the two constant polarization tests and randomly permuted the observations for 1000 iterations, then performed the same statistical test. The p-values were then determined by tallying the number of iterations where the fraction of permuted iterations exceeded that of the actual, unpermuted observations.

Likewise, to assess the fraction of channels exhibiting significant responses to constant or alternative polarization, we compared the magnitudes and latency of voltage extrema from trials with constant polarization to those with alternative polarization, employing the same procedure as for testing the channel selection to certain direction of polarization.

## Results

To study how squid photoreceptor responses to the movement of potential targets in caustics within shallow waters, we used linearly polarized flashes with constant intensity as a simple representation of the dominant parameters of underwater illumination (**Methods**). Across all experiments, we measured one retinal responses to 103 flashes across 8 trials, with each trial lasting 150 seconds. As a simplified representation of potential target movement in the flickering underwater environment, we rotated the stimulus polarization by 90º during the dark interval (46 flashes in 4 trials, **Methods**). As a comparison, we repeated a similar stimulus sequence, but did not rotate the polarization of the stimulus during the dark interval (57 flashes in 4 trials). On average, the light onsets were recorded every 10.9 ± 5.8 seconds (mean ± st.d.), with light-off intervals of 7.8 ± 4.9 seconds. Under these stimuli, extra-cellular voltages were recorded from 60 independent contacts to the anterior surface of the squid retina. Out of all recording channels, 57 of 60 (95%) exhibited a significant change in voltage extrema in response to the onset of illumination and analyzed further (see **Methods** for details).

### Retinal response to the onset of illumination

In the first analysis, we examined the characteristics of retinal response recorded extra-cellularly when there was a sudden illumination. Across all channels collectively, retinal responses were characterized by a fast initial maximum, followed by a subsequent minimum in voltages. Specifically, there was a discernible rise in voltage at 45 ms after the onset of light, reaching a peak of 463 μV at 55 ms, and exhibiting a rapid decline thereafter (**Figure 1C**). The width at full-width-half-maximum (FWHM) of the peak was 17 ms. Following the initial peak, we observed a negative potential, reaching a minimum of - 116 μV at 227 ms, and this feature had a FWHM of 225 ms.

This characteristic temporal pattern of retinal response was consistently observed across channels. However, we also observed significant variation in the magnitude of the extrema. The magnitudes of the maximum and the minimum exhibited a standard deviation of 193 and 50 μV across channels respectively, and the latency of these extrema showed standard deviations of 1 and 52 ms. We also found that these neural responses were not confined to specific contacts but were broadly distributed across the recording array (**Figure 1D**). To summarize, responses observed on different channels throughout the retina demonstrated similar shapes along the time axis, but variations in the specific amplitudes.

### Retinal response to specific orientation of polarizations

In a shallow underwater environment dominated by caustic flashes, a constant polarization amid fluctuations in light intensity suggests a reduced probability of encountering predators or companions, unlike situations involving changes in polarization (such as from the swimming motion of a silvery, specularly reflective predator). Next, we investigate the response of photoreceptors under illumination condition with constant light polarizations, which aims to extract features of the retinal response from steady background when exposed to flickering from caustics.

In this experiment, when averaged across the retina, neuronal responses were similar regardless of the fixed polarization orientation of the stimulus relative to the retina (**Figure 2A**). Specifically, the maxima were 404 ± 143 μV and 382 ± 189 μV for each of the two arbitrary polarizations tested, and 104 ± 37 μV and 111 ± 53 μV for the absolute values of the minima. Neither of these differences in extrema were significant across flashes (T-test, p > 0.35 for all features examined). Therefore, the retinal cells collectively exhibited a consistent response pattern irrespective of the specific polarization relative to the retina, indicating a robust neuronal response in processing visual stimuli under the constant polarizations.

**Figure 2.**
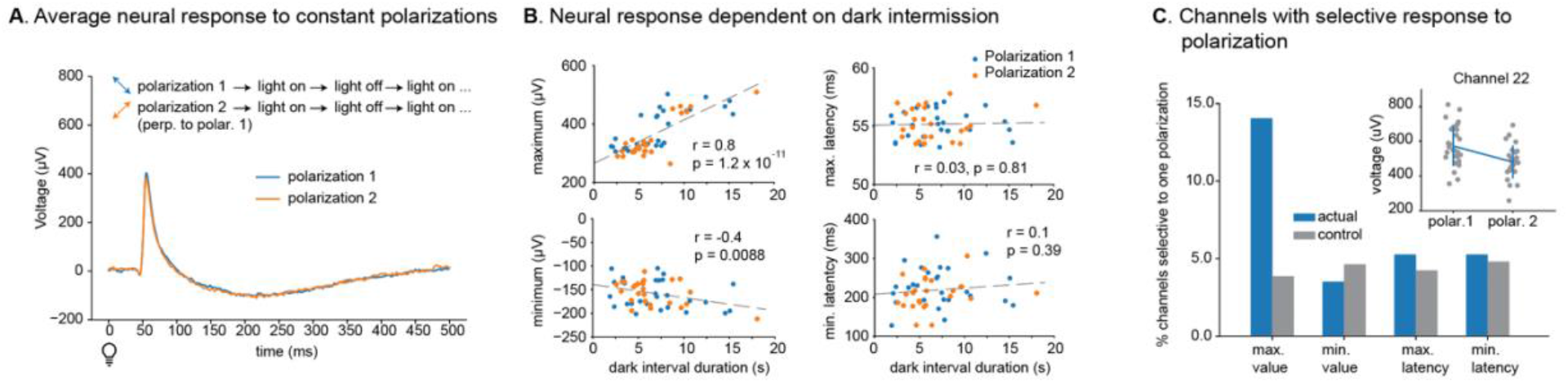
Retinal response to illumination onset with constant polarizations. **A**. Retinal response to two constant polarizations averaging over all recording channels. **B**. The extrema of the neural response depended on the duration of the dark interval before the onset of illumination, whereas the latency of the extrema did not demonstrate significant dependence on the dark duration. **C**. Across all channels recorded, 13% channels showed significant different maxima of neural response to illuminations with one polarization from the other, while the latencies of the extrema and the magnitude of refractory were not significantly different from random fluctuations. *Inset* showed an example channel with distinct peak voltages for each polarization. Gray dots represent the peak voltage responding to the onset of illumination.

However, at the level of individual channels, we observed that certain retinal responses were selective to the fixed polarizations. For example, in channel 22, the peak magnitude reached a voltage of 572 ± 114 μV for one polarization, which was significantly different from that of the other (480 ± 95 μV, p = 2 x 10^−3^, **Figure 2C**). Across all channels together, we observed 14% (8 out of 57) of channels with significant difference of the peak magnitude in one polarization compared to the other (permutation test, p = 0.057). The latencies of both extrema and the magnitude of the minimum were not significantly different between the two constant polarizations (permutation test, p > 0.3 for all features). Together, our results suggest that some regions of the retina were more responsive to specific polarization orientations than others given their outsize response to flashes with a fixed polarization orientation.

In addition to the response to the polarization, neural response also depended on the duration of the dark interval. Specifically, the magnitude of peak voltage was positively correlated to the light-off interval durations before the light onset (*r* = 0.8), which was significantly higher than random fluctuations of the signal (p = 1.2 x 10^−11^, **Figure 2B**). Similarly, the minima also became more extreme with an increase of the dark habituation durations (r = 0.4 and p = 9 x 10^−3^). However, the latencies of these two extrema did not depend on the dark durations prior to the light onset (p = 0.81 and 0.39, respectively).

### Retinal response enhanced to illumination with alternative polarizations

In shallow water environments, moving predators, especially those with specularly reflective skin, will alter light polarization relative to the background in a visual field. Hence, a discernible change in polarization between two illuminations at a given location serves as an indicative signal for the presence of moving objects requiring heightened vigilance. In this context, we conducted an analysis of retinal responses to light flashes with polarization orientation rotated by 90º during the inter-flash interval, and compared these responses to the experiment described above, in which the stimulus was polarized but did not rotate during the inter-flash interval.

Overall, the squid retina had a more active response when there was an alternation of polarizations between two light-on events, as compared to the same experiment with a constant polarization. In these trials with alternating polarization orientation, the average peak magnitude was 601 ± 185 μV, which was 1.5 times greater compared to the response to constant polarization (394 ± 167 μV). The absolute voltage of the minimum also increased from 107 ± 47 μV to 154 ± 47 μV, and the minimum latency increased from 216 ± 50 ms to 277 ± 51 ms. These results were robust after the consideration of the duration of the dark interval (T-test, p = 1 x 10^−5^ for the maxima and p = 0.01 for the minima, **Figure 3A**). Together, these findings suggest a significant increase in retinal neuron responsiveness when stimulated with alternating polarization orientation.

**Figure 3.**
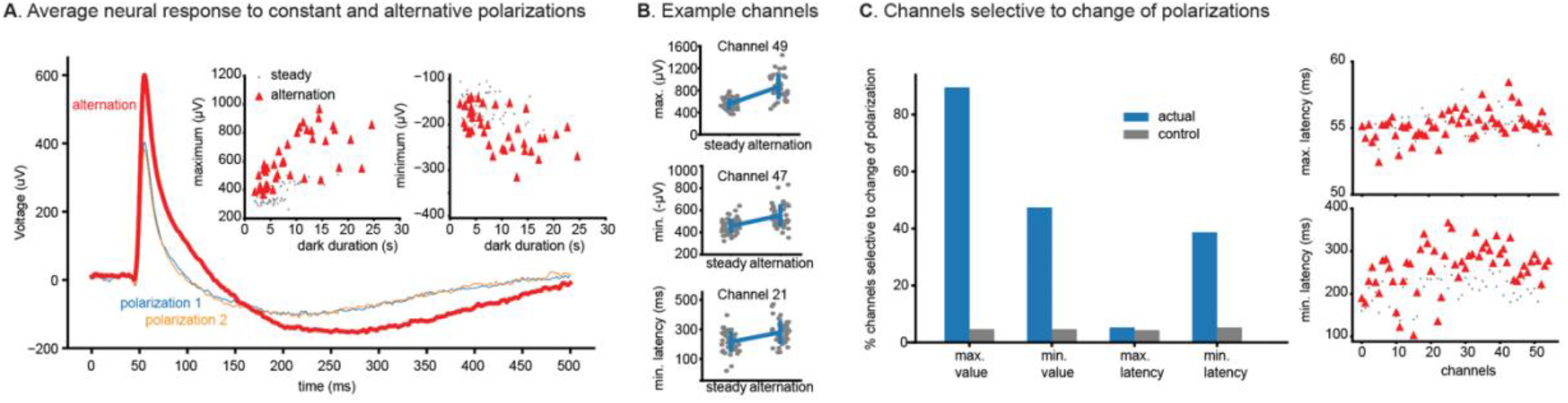
Retinal response to alternating orthogonal polarizations. **A**. Retinal responses were significantly increased when there was a change of polarization in between two illuminations. *Insets* show the voltage extrema of the neuronal responses across various dark durations for alternation and constant polarizations. **B**. Example channels illustrating the changes of both the magnitudes and latency for neuronal response to alternation polarizations. **C**. Most channels exhibited varying responses to changes in polarization compared to constant polarization, and these percentages were compared to random permutations of the events (*left*). The latencies of the extrema by channels are illustrated on the *right*.

When we compared individual channels across these two experiments, we observed that most of them exhibited enhanced responsiveness. For example, in channel 49, the peak magnitude increased from 553 ± 100 μV during constant polarization to 864 ± 225 μV in trials with rotating polarization (**Figure 3B**). Similarly, in channel 47, the magnitude of the minimum increased from 457 ± 63 μV to 554 ± 99 μV. In channel 21, the latency changed from 158 ± 168 ms to 281 ± 78 ms. Therefore, at the level of individual channels, retinal response showed enhanced magnitudes for the two extrema and a longer latency of the minimum when the illumination changed polarization.

Across all channels, there was a significantly higher percentage of channels with changing response to alternating polarization orientation (**Figure 3C**). Among the neural responses we examined, the most salient change was the magnitude of the peak: 89% channels showed a significant increase between trials where the orientation alternated between trials. This percentage was much higher than that of cells that responded selectively to one polarization orientation or the other (14%, **Figure 2C**). This result was also not due to random fluctuations in the recordings because a permutation of these peak magnitudes resulted in only 5% significant channels (p < 1 x 10^−3^). In addition, there were 47% of channels in which the value at the minimum decreased, and 39% of channels also showed an increase in the latency of the minimum (p < 1 x 10^−3^ for both). In comparison, there was no significant change in the latency of the maximum voltage, with only 5% of channels showing a selective response in the location of the peak (permutation test, p = 0.2). Together, these features indicate that retinal response in single channels significantly increased when there was a change of polarization orientation between flashes, and this response was not observed when the same retina exposed to flashes with constant polarization orientation.

## Discussion

To investigate how squid photoreceptors respond to changes in polarization indicative of moving objects in the caustic underwater environment, we utilized multi-electrode arrays to record extracellular retinal voltages under alternative polarization illumination. We showed the neural response to the onset of flashes recorded simultaneously across dozens of channels and their selectivity to certain direction of polarization. But more importantly, we found that retinal cells responded significantly more strongly to flashes with of alternating polarization orientation compared to flashes with constant polarization orientation. This result suggests that changes of polarization signals augmented neuronal response in the photoreceptor. These enhanced activities were observed across all channels recorded and were significantly higher than what can be explained by random fluctuation of neural recordings, suggesting a general and consistent neural response across retinal cells. Interestingly, we found that channels not selective to a specific polarization still increased their response to changes in polarization, which may potentially be attributed to the organization of the photoreceptors, where adjacent cells absorb perpendicularly polarized photons (*24*). Together, our findings indicate a clear neuronal response to the change of polarization, providing a preliminary explanation to the effective function of detecting moving object using polarization vision in a caustic underwater environment (*12*).

Our findings contribute to the broader understanding of polarization vision beyond shallow caustic water environments. Across various underwater ranges and environments, polarization vision has been demonstrated to play a crucial role in various advantages including navigation, predator and prey detection, gaze stability, and inter-subject communication (*8, 27-30*). These advantages are not limited to shallow waters, as marine animals with polarization vision in deeper water system also demonstrate an exceptional sensitivity to detect polarized moving objects, contrasting with fish, which rely primarily on chromatic vision (*31-33*). Our study provides a neural explanation that aligns well for such ability by revealing enhanced retinal photoreceptor responses in squid to changes in polarization, suggesting their role in utilizing polarization cues to facilitate vision in diverse water environments. Further, similar enhancements in neural responses have also been reported in crayfish (*34*), suggesting a convergence of neural mechanisms for polarization vision underwater, despite the considerable evolutionary distance between these species.

Although the focus of this work is not on the specific pathways of retinal cells in response to light (*35, 36*), our results do provide some insight into the underlying neuronal mechanism for polarized light detection. Using illuminations at constant intensity, we found the neuronal response included two extrema to polarized light, a maximum in voltage at 55 milliseconds followed by a refractory period with a minimum at 227 milliseconds. Although the exact voltages of the extrema varied across electrodes and changed for flashes with rotated polarizations, the latency of the maximum was remarkably constant, exhibiting little variance (s.t.d = 1 millisecond). This result indicates a universal neural mechanism characterized by precise and consistent temporal response. In contrast, the magnitude and the latency of the minimum showed a higher variance across channels and changed with the polarization condition of the flash, suggesting a different neuronal mechanism of the refractory period compared to the peak under the condition of changing stimulus. In addition, the magnitudes of the extrema were dependent on the dark duration before the light onset, suggesting a cellular adaptation that influences the magnitude of the retinal responses. Finally, extending previous findings of the tuning properties based observed on the action potentials (*17*), some channels exhibited increased sensitivity to specific polarizations, and displayed varying refractory periods, suggesting the complexity of the neural dynamics involved. Together, these findings illustrated detailed neural responses across channels and various polarization, with the specific features of responding voltages that are both preserved and modified in response to the light stimuli.

Collectively, these findings suggest a comprehensive and detailed collection of neural activities at the mesoscale level, potentially enabling cephalopods to efficiently detect moving objects in the dynamic, caustic underwater environment. This study had a limited sample size, such that it is difficult to draw more detailed conclusions about visual perceptions of polarization across individuals, though it is clear that such a neural response exists in squid. In future work, we will explore the mechanism of the enhanced response to polarization, i.e. whether it originates from isolated, individual cells or horizontal interactions between the cells. Nonetheless, our study provides a groundwork to begin investigating the neuronal basis of polarization vision using array-based neural recording technologies.

## Acknowledgements

The authors acknowledge support from NSF IOS-1343159 to A.M.S and a Packard Foundation fellowship to A.M.S. We are grateful to Vijay Balasubramaniam for sharing MEA resources with us. We acknowledge the use of ChatGPT for language refinement in this manuscript.

## References

1. J. A. Lock, J. H. Andrews, Optical caustics in natural phenomena. American journal of physics 60, 397–407 (1992).

2. Y. Swirski, Y. Y. Schechner, B. Herzberg, S. Negahdaripour, in 2009 IEEE 12th International Conference on Computer Vision. (IEEE, 2009), pp. 205–212.

3. Y. Swirski, Y. Y. Schechner, in The Nature of Light: Light in Nature IV. (SPIE, 2012), vol. 8480, pp. 69–76.

4. S. R. Matchette, I. C. Cuthill, N. E. Scott-Samuel, Concealment in a dynamic world: dappled light and caustics mask movement. Animal behaviour 143, 51–57 (2018).

5. S. R. Matchette, I. C. Cuthill, K. Cheney, N. J. Marshall, N. Scott-Samuel, Underwater caustics disrupt prey detection by a reef fish. Proceedings of the Royal Society B 287, 20192453 (2020).

6. D. Brewster, IX. On the laws which regulate the polarisation of light by reflexion from transparent bodies. By David Brewster, LL. DFRS Edin. and FSA Edin. In a letter addressed to Right Hon. Sir Joseph Banks, Bart. KBPR S. Philosophical Transactions of the Royal Society of London, 125–159 (1815).

7. A. Malins, The Reflectance of Silvery Fish Skins. (2009).

8. S. Johnsen, N. J. Marshall, E. A. Widder, Polarization sensitivity as a contrast enhancer in pelagic predators: lessons from in situ polarization imaging of transparent zooplankton. Philosophical Transactions of the Royal Society B: Biological Sciences 366, 655–670 (2011).

9. N. Shashar, R. T. Hanlon, A. d. Petz, Polarization vision helps detect transparent prey. Nature 393, 222–223 (1998).

10. L. M. Mathger, N. Shashar, R. T. Hanlon, Do cephalopods communicate using polarized light reflections from their skin? Journal of Experimental Biology 212, 2133–2140 (2009).

11. C. Drerup, M. J. How, J. E. Herbert-Read, Visual noise from caustic flicker does not affect the hunting success of cuttlefish. Animal Behaviour 202, 59–72 (2023).

12. S. V. Venables et al., Polarization vision mitigates visual noise from flickering light underwater. Science Advances 8, eabq2770 (2022).

13. N. Shashar, R. Hagan, J. G. Boal, R. T. Hanlon, Cuttlefish use polarization sensitivity in predation on silvery fish. Vision research 40, 71–75 (2000).

14. N. Shashar, C. Milbury, R. Hanlon, Polarization vision in cephalopods: neuroanatomical and behavioral features that illustrate aspects of form and function. Marine and Freshwater Behaviour and Physiology 35, 57–68 (2002).

15. C. M. Talbot, J. N. Marshall, The retinal topography of three species of coleoid cephalopod: significance for perception of polarized light. Philosophical Transactions of the Royal Society B: Biological Sciences 366, 724–733 (2011).

16. W. M. Saidel, N. Shashar, M. T. Schmolesky, R. T. Hanlon, Discriminative responses of squid (Loligo pealeii) photoreceptors to polarized light. Comparative Biochemistry and Physiology Part A: Molecular & Integrative Physiology 142, 340–346 (2005).

17. E. F. MacNichol Jr, W. E. Love, Electrical responses of the retinal nerve and optic ganglion of the squid. Science 132, 737–738 (1960).

18. J. R. Pungor, C. M. Niell, The neural basis of visual processing and behavior in cephalopods. Current Biology 33, R1106–R1118 (2023).

19. J. Cai, J. P. Townsend, T. C. Dodson, P. A. Heiney, A. M. Sweeney, Eye patches: Protein assembly of index-gradient squid lenses. Science 357, 564–569 (2017).

20. F. D. Hanke, A. Kelber, The eye of the common octopus (Octopus vulgaris). Frontiers in physiology 10, 1637 (2020).

21. M. F. Land, D.-E. Nilsson, Animal eyes. (OUP Oxford, 2012).

22. A. Fein, E. Z. Szuts, Photoreceptors: their role in vision. (CUP Archive, 1982), vol. 5.

23. H. R. Saibil, An ordered membrane-cytoskeleton network in squid photoreceptor microvilli. Journal of molecular biology 158, 435–456 (1982).

24. J. Z. Young, The retina of cephalopods and its degeneration after optic nerve section. Philosophical Transactions of the Royal Society of London. Series B, Biological Sciences 245, 1–18 (1962).

25. R. Williamson, M. Ichikawa, G. Matsumoto, Neuronal circuits in cephalopod vision. Netherlands journal of zoology, (1994).

26. W. M. Saidel, J. Y. Lettvin, E. F. MacNichol Jr, Processing of polarized light by squid photoreceptors. Nature 304, 534–536 (1983).

27. N. Shashar et al., Underwater linear polarization: physical limitations to biological functions. Philosophical Transactions of the Royal Society B: Biological Sciences 366, 649–654 (2011).

28. I. M. Daly et al., Dynamic polarization vision in mantis shrimps. Nature communications 7, 12140 (2016).

29. L. Cartron et al., Polarization vision can improve object detection in turbid waters by cuttlefish. Journal of Experimental Marine Biology and Ecology 447, 80–85 (2013).

30. G. Horváth, A. Lerner, N. Shashar, Polarized light and polarization vision in animal sciences. (Springer, 2014), vol. 2.

31. L. Nahmad-Rohen, M. Vorobyev, Spatial contrast sensitivity to polarization and luminance in octopus. Frontiers in Physiology 11, 379 (2020).

32. J. C. Tuthill, S. Johnsen, Polarization sensitivity in the red swamp crayfish Procambarus clarkii enhances the detection of moving transparent objects. Journal of experimental biology 209, 1612–1616 (2006).

33. V. Pignatelli et al., Behavioural relevance of polarization sensitivity as a target detection mechanism in cephalopods and fishes. Philosophical Transactions of the Royal Society B: Biological Sciences 366, 734–741 (2011).

34. R. M. Glantz, Polarization vision in crayfish motion detectors. Journal of Comparative Physiology A 194, 565–575 (2008).

35. R. Clark, G. Duncan, Two components of extracellularly-recorded photoreceptor potentials in the cephalopod retina: differential effects of Na+, K+ and Ca 2+. Biophysics of structure and mechanism 4, 263–300 (1978).

36. A. Chrachri, L. Nelson, R. Williamson, Whole-cell recording of light-evoked photoreceptor responses in a slice preparation of the cuttlefish retina. Visual neuroscience 22, 359–370 (2005).

